# FosB/ΔFosB activation in mast cells regulates gene expression to modulate allergic inflammation in male mice

**DOI:** 10.1101/2024.05.06.592755

**Authors:** Natalia Duque-Wilckens, Dimitri Joseph, Meesum Syed, Brianna Smith, Nidia Maradiaga, Szu-Ying Yeh, Vidhula Srinivasan, Fabiola Sotomayor, Kait Durga, Eric Nestler, Adam J Moesers, A.J. Robison

## Abstract

Mast cells are innate immune cells that regulate physiological processes by releasing pre-stored and newly synthesized mediators in response to allergens, infection, and other stimuli. Dysregulated mast cell activity can lead to multisystemic pathologies, but the underlying regulatory mechanisms remain poorly understood. We found that FOSB and ΔFOSB, transcription factors encoded by the *FosB* gene, are robustly expressed in mast cells following IgE-antigen stimulation, suggesting a role in modulating stimulus-induced mast cell functions. Using phenotypic, gene binding, and gene expression analyses in wild-type and mast cell-specific *FosB* knockout male mice, we demonstrate that FOSB/ΔFOSB modulates mast cell functions by limiting reactivity to allergen-like stimuli both *in vitro* and *in vivo*. These effects seem to be mediated, at least in part, by FOSB/ΔFOSB-driven enhanced expression of DUSP4, a dual-specificity phosphatase that attenuates MAPK signaling. These findings highlight FOSB/ΔFOSB as critical regulators of mast cell activity and potential targets for therapeutic intervention.

**Graphical Abstract:** 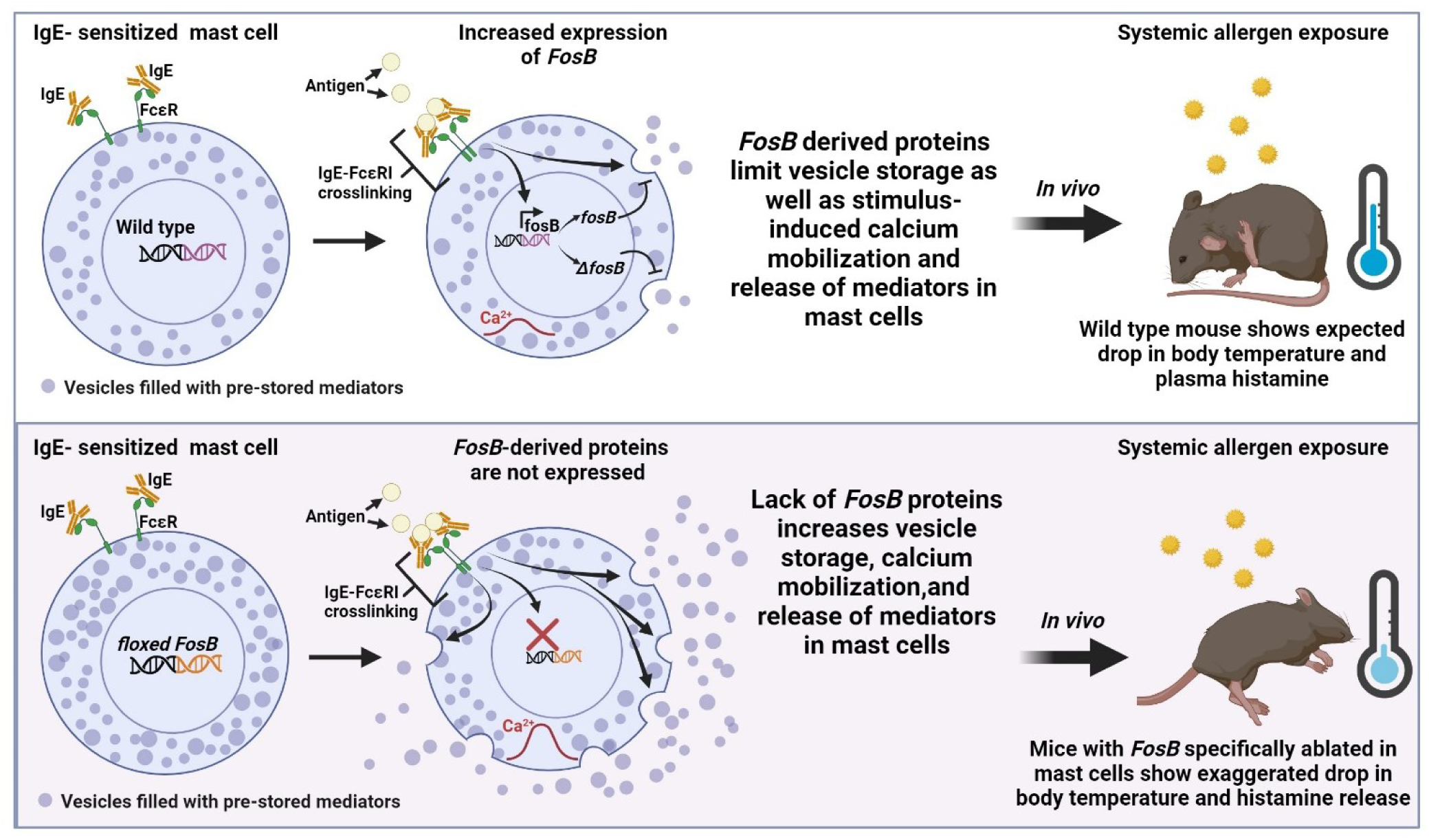

## Introduction

Mast cells are highly conserved immune cells that play a central role in diverse physiological processes. They are distributed throughout tissues, particularly at barrier surfaces such as the skin and near blood and lymphatic vessels^1,2^ and possess an extensive repertoire of receptors that enable them to detect and rapidly respond to a wide range of stimuli. These include pathogens and cellular damage^3^, environmental toxins^4,5^, immune cells and signaling molecules^6–10^, neurotransmitters and neuromodulators^11,12^, as well as hormones involved in stress^13,14^, metabolism^15,16^, sexual development^17,18^, and circadian rhythms^19^. Further, upon activation, mast cells can selectively release a variety of pre-stored and newly synthesized mediators, such as cytokines and chemokines^20^, lipid mediators (e.g., prostaglandins and leukotrienes)^21^, neurotransmitters (e.g., serotonin^22^, histamine^23^, dopamine^24^), and growth factors (e.g. VEGF^25^ and NGF^26^). Consequently, mast cells are involved in initiating and orchestrating innate and adaptive immune responses^27,28^, driving tissue development^29,30^ and repair^31,32^, influencing metabolic processes^33,34^, and modulating neuronal functions^35,36^ and behavior^37–39^, among many others^40,41^.

Given their extensive roles, it is unsurprising that disruptions in mast cell function can contribute to the pathology of numerous inflammatory diseases. For example, mast cell dysfunctions are implicated in allergic disorders^42^, migraines^43,44^, inflammatory bowel disease^45,46^, multiple sclerosis^47^, rheumatoid arthritis^48^, and chronic neuroinflammatory conditions, such as Alzheimer’s disease^49^. Thus, understanding the mechanisms that regulate mast cell gene expression and activity is crucial for developing targeted strategies to enhance their protective functions while minimizing their harmful effects across systems.

In an RNA-seq screen investigating gene regulation in mouse bone marrow-derived mast cells (BMMCs) following antigen-induced IgE-FcεRI crosslinking—a key mechanism of mast cell activation during allergic inflammation^50^—we identified *FosB* as one of the most robustly upregulated genes. Notably, *FosB* was also among the most strongly upregulated genes in a separate study where BMMCs were stimulated with glutamate^12^, a neurotransmitter implicated in neuron-mast cell communication during tissue injury. Recent work by Tauber et al.^1^, integrating whole-tissue imaging and single-cell RNA sequencing, further revealed that *FosB* is expressed across multiple human mast cell populations, including those in the skin, bladder, lymph nodes, and vasculature —highlighting its potential as a conserved regulator of mast cell functions.

*FosB* is well-characterized in neurons as an immediate early gene encoding two primary proteins, FOSB and ΔFOSB, via alternative splicing. Both proteins dimerize with Jun family members to form activator protein-1 (AP-1) transcription factor complexes, regulating gene expression across diverse signaling pathways^51–54^. In neurons, FOSB is rapidly degraded after induction, whereas ΔFOSB is remarkably stable, allowing it to modulate gene expression long-term. This sustained activity underlies chronic stress adaptation^51,55,56^, memory formation^57^, and responses to drugs of abuse^58^. However, whether FOSB and ΔFOSB serve similar gene-regulatory roles in mast cells has not been explored.

To begin addressing this gap, we first validated our RNA-seq findings and confirmed that both FOSB and ΔFOSB isoforms are markedly upregulated by IgE-antigen–mediated activation in BMMCs at both the mRNA and protein levels. Next, we employed Cleavage Under Targets and Release Using Nuclease (CUT&RUN) to map genome-wide changes in FOSB/ΔFOSB DNA occupancy in IgE-sensitized BMMCs before and after antigen stimulation. We found that FOSB/ΔFOSB in IgE-sensitized BMMCs bind to multiple genes associated with mast cell proliferation, mediator storage, and activation, and that antigen exposure further enhances binding at genes associated with calcium mobilization and degranulation.

To determine the functional relevance of these DNA binding patterns, we crossed the mast cell-specific Cre reporter, *Mcpt5-Cre*^59^, with the Cre-dependent *floxed FosB*^60^ mouse lines to ablate *FosB* specifically in Mcpt5-expressing mast cells (MC^FosB⁻^) —a widely distributed population found in connective and other tissues, including the peritoneal cavity, mesentery, dura mater, skin^1,61,62^. We found that, in response to IgE-antigen stimulations, MC^FosB⁻^ BMMCs show increased histamine content, increased sensitivity to activation, and increased release of mediators compared to wild type (WT) BMMCs. Moreover, MC^FosB⁻^ male mice show exaggerated hypothermic responses, tissue-resident mast cell degranulation, and plasma release of histamine *in vivo*, suggesting an important role for FOSB/ΔFOSB in limiting mast cell reactivity to allergen-like stimuli.

Finally, we show that although MC^FosB⁻^ and WT BMMCs exhibit comparable IL-6 secretion in response to lipopolysaccharide (LPS) stimulation *in vitro*, MC^FosB⁻^ mice display exacerbated hypothermic responses and elevated plasma IL-6 levels following LPS challenge *in vivo*. These findings suggest that FOSB/ΔFOSB in mast cells modulate downstream activation of other immune cells that drive LPS-induced sickness and inflammation.

In sum, this study reveals a critical role for *FosB*-derived proteins in limiting mast cell responses to allergens, with potential implications for inflammatory regulation beyond allergy. Fully elucidating how FOSB/ΔFOSB regulate these processes could inform novel therapeutic strategies for mitigating allergic and inflammatory diseases. While this study does not distinguish between the specific roles of FOSB and ΔFOSB, the stability of ΔFOSB presents intriguing possibilities for long-term modulation of mast cell function, highlighting a compelling avenue for future research.

## Results

### 1. *FosB* Splice Variants FOSB and ΔFOSB Are Upregulated After IgE-Antigen Stimulation in BMMCs and Exhibit Stimulation-Dependent Gene Binding

Building on our initial RNA screening results, which identified *FosB* as one of the most upregulated genes following IgE-FcεRI crosslinking, we first sought to validate these findings using qPCR and Western blot in male-derived BMMCs. BMMCs were sensitized overnight with 0.8 µg/ml IgE anti-2,4-dinitrophenyl (DNP) and subsequently challenged with vehicle or 15.5 ng/ml DNP (**Fig. 1A**), as described previously^63,64^. DNP exposure significantly increased the expression of both *fosB* and *ΔfosB* mRNAs (**Fig 1B&C**), along with a fourfold increase in FOSB and ΔFOSB protein levels **(Fig 1 D-F),** suggesting a role in regulating the transcriptional response of BMMCs to IgE-antigen stimulation. Interestingly, while *fosB* and *ΔfosB* mRNA levels were minimal in vehicle-treated IgE-sensitized BMMCs, FOSB and ΔFOSB proteins were readily detectable, suggesting that these proteins are either constitutively maintained in BMMCs or persist long after (>12h) an IgE-induced response.

**Fig. 1.**
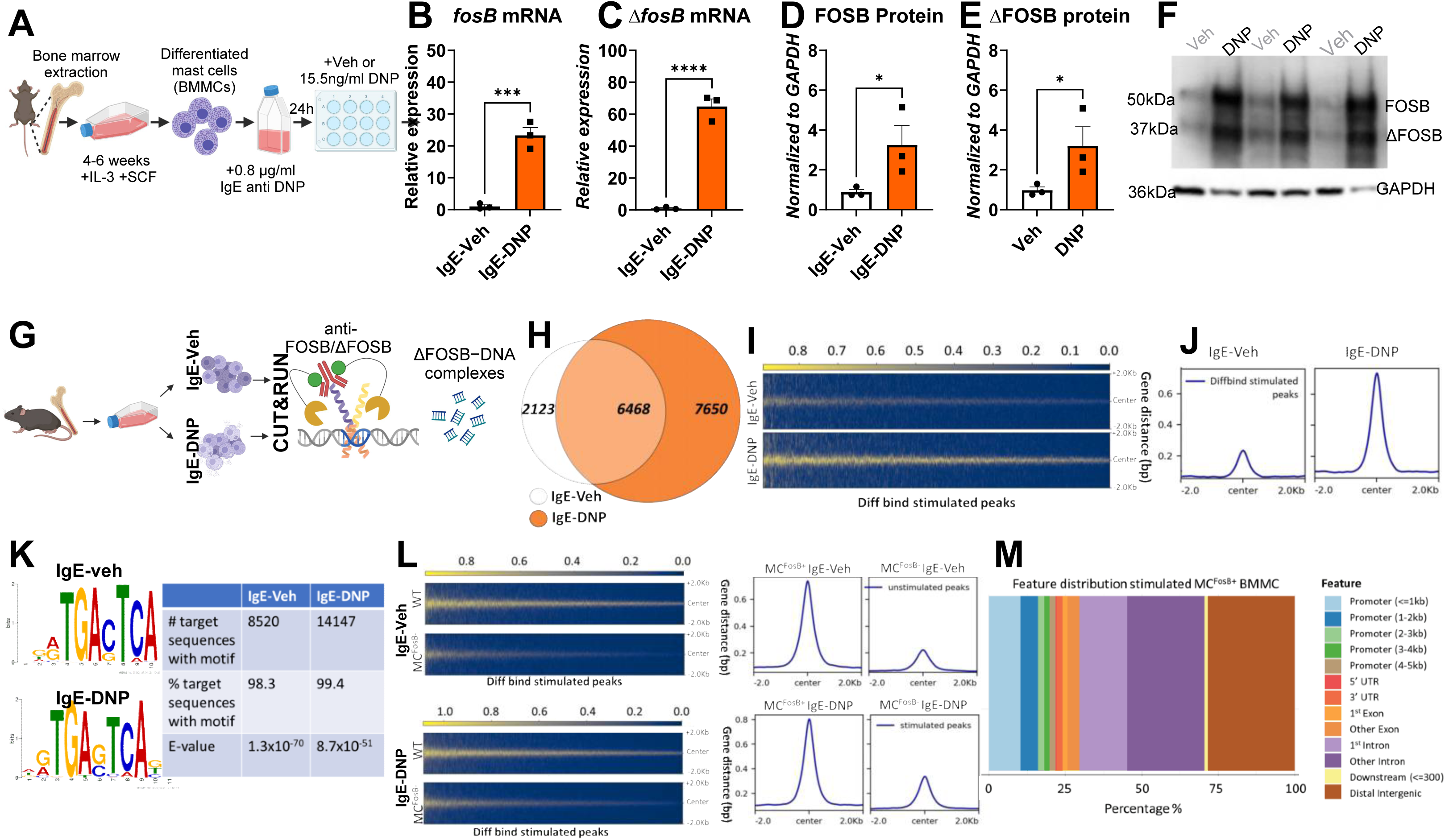
***FosB* Splice Variants FOSB and ΔFOSB Are Upregulated After IgE-Antigen Stimulation in BMMCs and Exhibit Stimulation-Dependent Gene Binding A.** Timeline showing BMMC isolation, culture, and DNP-mediated IgE-FcεRI crosslinking. **B,C.** Relative expression of *fosb* (n=3/group, one-tailed T-test t_(4)_=8.96, p=0.0004) and Δ*fosb* (n=3/group one-tailed t-test t_(4)_=13.50, p<0.0001) 1h after veh/DNP. **D,E**. Expression of FOSB (n=3/group, one-tailed T-test t_(4)_=2.411, p=0.004) and ΔFOSB (n=3/group, one-tailed T-test t_(4)_=2.28, p=0.04) at baseline and after DNP stimulation in IgE-sensitized BMMCs 4h after veh/DNP. **F.** Western blot showing expression of FOSB (50kDa) and ΔFOSB (37 kDa). **G.** Schematic of CUT&RUN experimental workflow in IgE-sensitized BMMCs using anti-FOSB antibody. **H.** FOSB/ΔFOSB**-**binding peak comparison between stimulated and unstimulated WT BMMCs (3 unstim and stimulated BMMC replicates derived from 3 different adult 8 week old mice). **I.** Heatmaps and average profile plots showing normalized FOSB/ΔFOSB CUT&RUN signal ±2 kb from peak centers of differentially bound regions identified in stimulated BMMCs. Signal intensity is higher in DNP-stimulated BMMCs (left) compared to unstimulated controls (right), indicating stimulus-induced binding. **J.** Average CUT&RUN signal intensity (±2 kb from peak center) across all reproducible FOSB/ΔFOSB peaks in IgE-vehicle (top) and IgE-DNP (bottom) BMMCs. Antigen stimulation increased both the magnitude and sharpness of FOSB/ΔFOSB binding. **K.** Top enriched DNA motifs from FOSB/ΔFOSB-bound regions in unstimulated (top) and stimulated (bottom) BMMCs. Both conditions showed significant enrichment of the canonical AP-1 binding motif. **L.** CUT&RUN signal intensity comparison between wild type (WT) and *FosB*-deficient (MC^FosB⁻^) BMMCs at FOSB/ΔFOSB peak regions identified in IgE-veh (top) or IgE-DNP (bottom) conditions. Heatmaps show normalized read density ±2 kb around peak centers. Line plots depict average signal intensity across the same regions. The minimal peak enrichment in MC^FosB⁻^ BMMCs confirms the specificity of the CUT&RUN signal and validates successful ablation of FOSB/ΔFOSB. **M.** Genomic annotation of differentially bound FOSB/ΔFOSB peaks in IgE-DNP BMMCs. The stacked bar chart shows the percentage of peaks mapping to each genomic feature, including promoters, exons, introns, UTRs, downstream elements, and distal intergenic regions. The majority of stimulus-induced FOSB/ΔFOSB binding occurs within intronic and intergenic regions, suggesting regulatory activity beyond canonical promoter sites.

To explore whether FOSB and ΔFOSB directly bind to DNA in mast cells and identify the specific genes they target in the context of IgE-mediated activation, we conducted Cleavage Under Targets and Release Using Nuclease (CUT&RUN^65^) profiling in IgE-sensitized BMMCs exposed to vehicle or DNP (**Fig. 1G**). The anti-FOSB antibody used in this study does not distinguish between FOSB and ΔFOSB isoforms; therefore, we will collectively refer to these proteins in this context as FOSB/ΔFOSB.

We identified 8,664 FOSB/ΔFOSB peaks unique to IgE-vehicle and 14,228 peaks unique to IgE-DNP BMMCs, with 6,468 peaks shared across both conditions (**Fig. 1H**). This suggests a widespread redistribution of FOSB/ΔFOSB binding sites in response to IgE-antigen crosslinking. The majority of FOSB/ΔFOSB binding events occurred at conserved genomic regions, as indicated by the consistent peak distribution across conditions (**Fig. 1I&J**). More than 90% of FOSB/ΔFOSB-bound regions contained AP-1 motifs, the canonical response elements for FOS family proteins complexed with JUN family proteins, replicating previous findings in mouse neurons^66^ **(Fig. 1K)**. To confirm that these peaks represent *bona fide* FOSB/ΔFOSB binding sites, we applied the same methodology to BMMCs from mice with mast cell-specific *FosB* ablation (MC^FosB-^, see below) and found that peak coverage was dramatically reduced compared to wild type (WT) BMMCs (**Fig. 1L**). To further confirm specificity, we used Pearson correlation coefficient to compare the coverage of FOSB/ΔFOSB at peaks in IgE-veh and IgE-DNP WT and MC^FosB-^ BMMCs (**Supplementary Fig. 1A**). WT IgE veh and IgE-DNP BMMCs showed moderate-to-high correlation, suggesting that some ΔFOSB binding sites are present at baseline and become further enriched upon stimulation. In contrast, IgE-veh and IgE-DNP MC^FosB-^ vs. WT BMMCs showed a much lower correlation, indicating that FOSB/ΔFOSB-dependent binding is largely absent in knockout cells across conditions.

Next, we annotated the genomic regions bound by FOSB/ΔFOSB to determine their locations relative to the mouse reference genome (mm10). Approximately 15% of peaks were found within putative promoter regions (≤1000 bp upstream or ≤2000 bp downstream of known transcription start sites), 28% were intergenic, and nearly half were within gene bodies (**Fig. 1M**). Notably, this enrichment of FOSB/ΔFOSB binding outside promoter regions was recently reported in mouse neurons^66^, suggesting a broader regulatory role beyond direct promoter activation. To investigate the potential functional relevance of these FOSB/ΔFOSB-bound regions, we used HOMER^67^ to annotate each peak with its nearest gene, linking DNA-binding events to possible gene regulatory functions. This yielded two gene lists representing FOSB/ΔFOSB binding sites in unstimulated (**Supplementary Table 1**) and stimulated (**Supplementary Table 2**) WT BMMCs. To identify DNA regions differentially targeted by FOSB/ΔFOSB after antigen exposure, we performed differential analysis using the DiffBind package (v3.2.7)^68^, considering regions with an absolute log2 fold change of at least 1 and an FDR less than 0.05 as significantly different. This approach identified over 4,000 peak regions with substantially higher FOSB/ΔFOSB coverage following stimulation (**Supplementary Table 3**). To gain insight into the potential biological functions of FOSB/ΔFOSB-bound regions, we next performed Gene Ontology (GO) molecular enrichment analyses (http://geneontology.org/).

GO analysis of FOSB/ΔFOSB-bound genes in IgE-Veh BMMCs identified several functional categories enriched in pathways relevant to histamine storage, mast cell growth, and cytoskeleton remodeling, including amine binding (GO: 0043176)^69^, platelet-derived growth factor receptor binding^70^ (GO:0005161) and actin filament binding^71^ (GO:0051015). Additionally, several terms associated with IgE-mediated mast cell activity and proliferation^7,72–74^ were enriched, including those related to MAPK signaling (MAP kinase tyrosine/serine/threonine phosphatase activity (GO:0017017), mitogen-activated protein kinase binding (GO:0051019)), as well as cytokine signaling pathways such as interleukin-6 receptor binding (GO:0005138) and tumor necrosis factor receptor binding (GO:0005164) (**Fig. 2A for list of selected terms, for full list see Supplementary Table 4**).

**Fig. 2.**
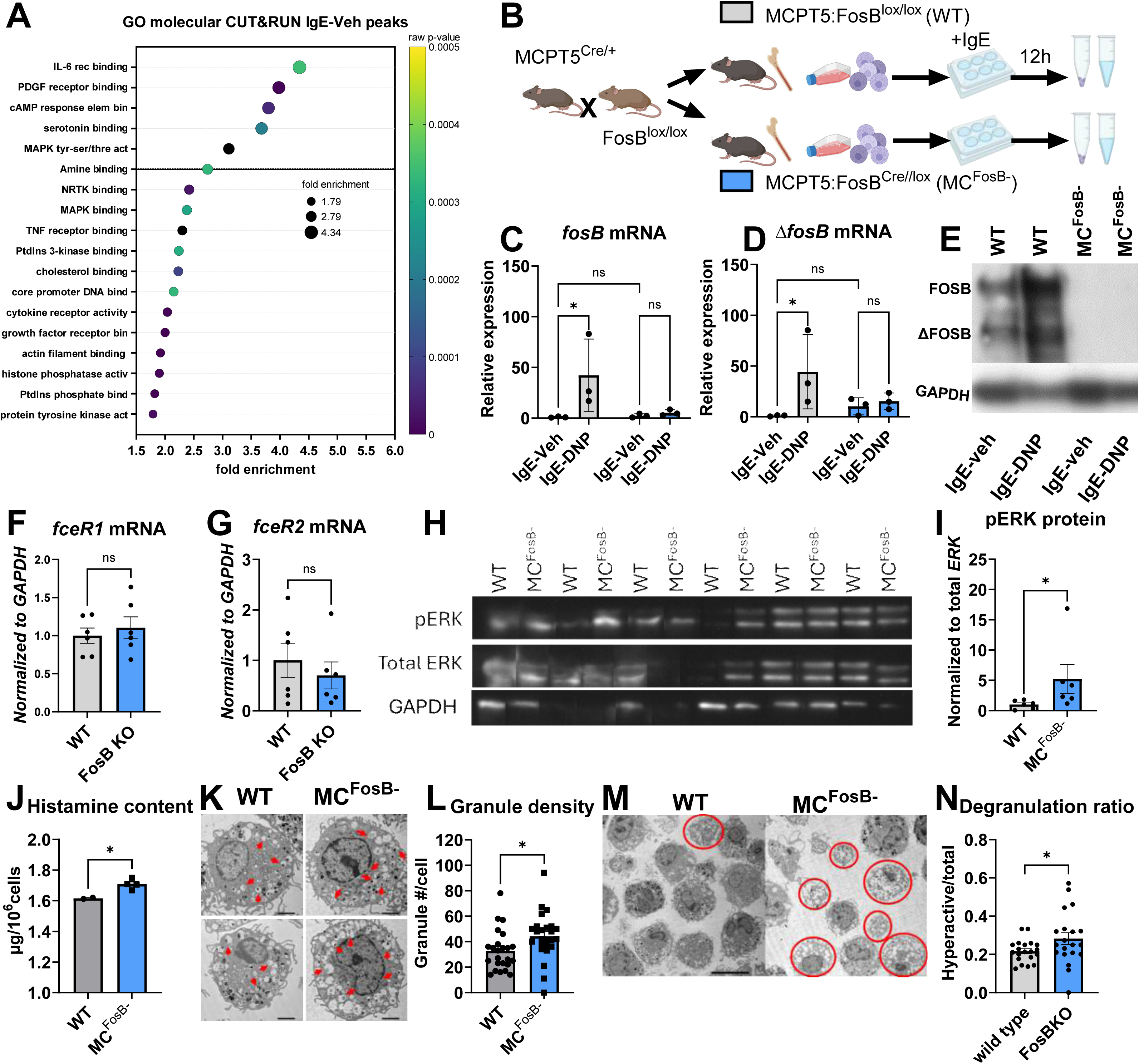
Inhibition of FosB gene expression in BMMCs enhances storage capacity and increases sensitivity and histamine release in response to IgE-mediated activation via pathways downstream of FcεRI. **A.** Gene ontology (GO) molecular function analysis of FOSB/ΔFOSB-bound peaks in IgE–vehicle–treated BMMCs reveals enrichment of functional terms related to signaling (e.g., MAPK binding, IL-6 receptor binding), mediator storage (e.g., amine binding, serotonin binding), and mast cell activation. **B.** Schematic of the experimental timeline for deriving BMMCs from WT and MC^FosB⁻^ mice. **C.** Relative expression of *fosb* (n=3/group, two-way ANOVA treatment F_(1, 8)_ = 4.5, p=0.06, Fisher’s LSD WT IgE-Veh vs. IgE-DNP p=0.02) mRNA after DNP stimulation in IgE-sensitized WT and MC^FosB-^ BMMCs. **D.** Relative expression of *Δfosb* (n=3/group, two-way ANOVA treatment F_(1, 8)_ = 4.8, p=0.06, Fisher’s LSD WT IgE-Veh vs. IgE-DNP p=0.02) **E.** Representative Western blot showing increased FOSB (50 kDa) and ΔFOSB (37 kDa) protein in WT BMMCs following DNP stimulation, with no expression detected in MC^FosB⁻^ cells. **F, G.** Expression of *Fcer1a* and *Fcer2a* mRNA at baseline in WT and MC^FosB⁻^ BMMCs (n=6/group); no significant genotype differences. **H, I.** Expression of phosphorylated ERK (pERK) normalized to total ERK in IgE-sensitized WT and MC^FosB⁻^ BMMCs (n=6 flasks per group, from 3 animals; Mann–Whitney exact *p*=0.015), showing enhanced ERK activation in MC^FosB⁻^ cells**. J.** Histamine content at baseline is elevated in MC^FosB⁻^compared to WT BMMCs (WT n=2, MC^FosB⁻^ n=4; two-tailed t-test, t_(4)_=3.642, *p*=0.02). **K.** Representative electron microscopy images illustrating granule density in WT and MC^FosB⁻^ BMMCs; empty granules indicated by red arrows. **L.** Quantification of granule number per cell from EM images (n=24 cells/genotype; two-tailed t-test, t_(47)_ =2.3, *p*=0.02). **M**. EM images used to assess degranulation ratio in BMMCs; cells with extensive empty granules (circled in red) indicate hyperactivity. **N.** Degranulation ratio is significantly increased in MC^FosB⁻^ BMMCs relative to WT (n=20 images/group; Mann– Whitney exact *p*=0.04).

To further investigate how FOSB/ΔFOSB activity is dynamically regulated in response to antigen exposure, we next performed GO analysis on the differentially bound gene list, representing genomic regions with significant changes in FOSB/ΔFOSB occupancy between IgE-Veh and IgE-DNP conditions. This analysis revealed key molecular functions linked to mast cell activation and degranulation. These included AP-3 adaptor complex binding, which regulates secretory granule trafficking^75^; calcium mobilization (e.g., calcium, potassium:sodium antiporter activity), essential for mast cell degranulation^76^; and JUN kinase kinase activity, a key signaling pathway driving mast cell inflammatory responses through cytokine production^77–79^ (**Fig. 3A for list of selected terms, for full list see Supplementary Table 5**).

**Fig. 3.**
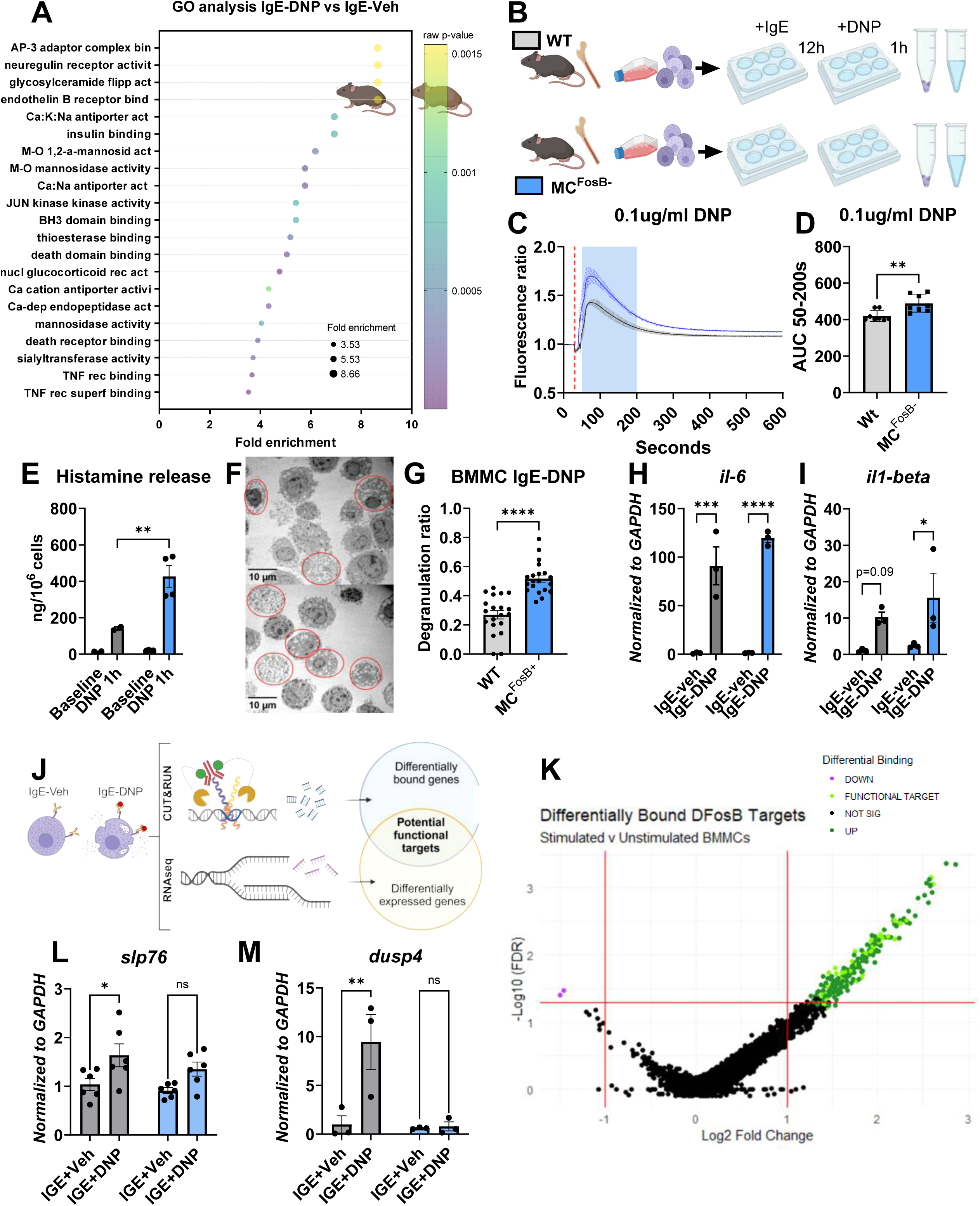
Inhibition of *FosB* gene expression in BMMCs exacerbates activation and histamine release in response to IgE-DNP stimulation.A. Gene ontology (GO) molecular function analysis of FOSB/ΔFOSB-bound peaks in IgE– vehicle–treated BMMCs identifies enriched terms associated with signaling and immune regulation (e.g., TNF receptor binding, calcium-dependent endopeptidase activity, and JUN kinase kinase activity). **B**. Experimental timeline of BMMC isolation, sensitization, and IgE-DNP stimulation. **C.** Calcium mobilization in WT and MC^FosB-^ BMMCs in response to IgE-DNP stimulation (n=8/group, repeated measures two-way ANOVA genotype x time interaction effect F_(999, 13986)_=3.597 p <0.0001. **D.** Area under the curve (AUC) for calcium flux between seconds 50–200 post-stimulation shows enhanced response in MCFosB⁻ BMMCs (one-tailed T-test t_(14)_=3.5, p=0.002, *r*_Yl_=0.65).**E.** Histamine release at baseline and 1h after DNP stimulation in IgE-sensitized WT and MC^FosB-^ BMMCs (n=2 WT, 4 MC^FosB-^, two-way ANOVA interaction F_(1, 8)_ = 9.9, p=0.01, Fisher’s LSD DNP WT vs MC^FosB-^ p=0.002, *r*_Yl_=0.92; vehicle WT vs MC^FosB-^ *ns*). **F**. Representative electron microscopy images of BMMCs show increased granule depletion in MC^FosB⁻^ cells after DNP stimulation (empty granules circled in red). **G**. Degranulation ratio is significantly increased in MC^FosB⁻^ BMMCs relative to WT (n=20 images/group; Mann–Whitney exact *p*=0.04). **H.** Relative expression of *il-6* (n=3/group, two-way ANOVA treatment F_(1, 8)_ = 10.5, p=0.01, Fisher’s LSD WT veh-DNP p=0.002, *r*_Yl_=0.88; MC^Fosb-^ veh-DNP p=<0.0001, *r*_Yl_=1) and **I.** *il1-beta* (n=3/group, two-way ANOVA treatment F_(1, 8)_ = 108, p<0.0001, Fisher’s LSD WT veh-DNP *ns*, MC^Fosb-^ veh-DNP p=0.03) mRNA 1h after DNP stimulation in IgE-sensitized WT and MC^FosB-^ BMMCs. **J.** Schematic of the data integration strategy combining CUT&RUN (differential FOSB/ΔFOSB binding) and RNA-seq (differential gene expression) to identify functional targets. **K.** Schematic of the data integration strategy combining CUT&RUN (differential FOSB/ΔFOSB binding) and RNA-seq (differential gene expression) to identify functional targets. **L.** Relative expression of *lcp2/slp-76* (n=6/group two-way ANOVA treatment F_(1,20)_ =11, p=0.003; Fisher’s LSD WT IgE veh vs WT IgE DNP p=0.01, MC^FosB-^ IgE veh vs MC^FosB-^ IgE DNP ns) and **M**. *dusp4* (n=3/group two-way ANOVA genotype F_(1,8)_ =9, p=0.02; Fisher’s LSD WT IgE veh vs WT IgE DNP ns, MC^FosB-^ IgE veh vs MC^FosB-^ IgE DNP p=0.008) mRNA after DNP stimulation in IgE-sensitized WT and MC^FosB-^ BMMCs.

Next, to determine the functional impact of FOSB/ΔFOSB on mast cells, we generated a novel mouse model with targeted *FosB* gene ablation specifically in mast cells (MC^FosB-^) by crossing the *Mcpt5-Cre*^80^ and *floxed FosB*^60^ mouse lines.

### 2. Inhibition of *FosB* gene expression in BMMCs enhances storage capacity and increases sensitivity and histamine release in response to IgE-mediated activation via pathways downstream of FcεRI

We derived BMMCs from adult male WT and MC^FosB⁻^ mice (**Fig. 2B**) and confirmed mast cell purity in both groups by co-expression of c-Kit and FcεRI (**Supplementary Fig. 1B**). To assess the effectiveness of *FosB* gene inhibition, we first measured *FosB* expression in IgE-sensitized BMMCs. In IgE-Veh-treated BMMCs, *fosB* (**Fig. 2C**) and *ΔfosB* mRNA (**Fig. 2D**) expression was minimal in both WT and MC^FosB⁻^ BMMCs. Intriguingly, although not statistically significant, BMMCs from MC^FosB⁻^ mice exhibited a baseline *ΔfosB* expression that appeared higher than that of WT BMMCs. This could result from PCR primers detecting residual mRNA fragments still expressed after recombination of the floxed gene. Alternatively, it may reflect incomplete Cre recombinase activity resulting from some BMMCs not expressing MCPT5. In contrast, following IgE-DNP stimulation, *fosB* and *ΔfosB* levels significantly increased in WT BMMCs, replicating initial findings, but remained unchanged in MC^FosB⁻^ BMMCs. Of greater significance for biological results, western blot analyses showed that while FOSB and ΔFOSB proteins were expressed in WT BMMCs both at baseline and to an even greater extent following stimulation, and no detectable expression of either protein was observed in MC^FosB⁻^ BMMCs (**Fig. 2E**), indicating successful functional elimination of *FosB* proteins.

Next, we investigated the impact of *FosB* knockout on IgE-sensitized BMMCs (**Fig. 2F**). We found no differences in the mRNA expression of the high-affinity IgE receptor FcεRI (**Fig. 2G**) or the low-affinity IgE receptor FcεRII (**Fig. 2H**). However, MC^FosB⁻^ BMMCs exhibited increased phosphorylation of ERK (pERK) (**Fig. 2I**), a key signaling event downstream of FcεRI activation that regulates degranulation and cytokine production^81^. Additionally, MC^FosB⁻^ showed a higher total content of histamine - one of the predominant bioactive amines stored in mast cell granules and extensively released in response to IgE-FcεRI crosslinking^23^-compared to WT BMMCs (**Fig. 2J**). Lastly, transmission electron microscopy confirmed that MC^FosB⁻^ BMMCs retained a typical mast cell phenotype, characterized by a large nucleus and multiple electron-dense granules^82,83^. However, compared to WT, MC^FosB⁻^ BMMCs displayed a significantly greater number of granules (**Figs. 2K,L**), indicating enhanced storage capacity, along with an increased degranulation ratio (**Fig. 2M,N**), suggesting heightened baseline activation. These results are consistent with the GO terms associated with FOSB/ΔFOSB binding in IgE-sensitized BMMCs associated with mast cell growth, histamine storage, and MAPK signaling (**Fig. 2A**).

We next examined functional response to IgE-DNP stimulation. Based on the marked increase in FOSB and ΔFOSB expression following DNP treatment, as well as the identification of FOSB/ΔFOSB DNA binding sites in key regulatory regions revealed by CUT&RUN analysis (**Fig. 3A**), we aimed to determine whether the loss of *FosB* alters activation markers and mediator release upon IgE-DNP challenge (**Fig. 3B**). We found that, in MC^FosB-^ BMMCs compared to WT BMMCs, DNP stimulation induced significantly higher calcium mobilization (**Figs. 3C&D**) and histamine release (**Fig. 3E**). Consistent with these findings, electron microscopy analyses revealed a significantly higher degranulation ratio in MC^FosB-^ BMMCs compared to WTs (**Figs. 3F&G**). However, IgE-DNP-induced *il-6* (**Fig. 3H**) and *il1-beta* (**Fig. 3I**) expression did not differ between genotypes.

Together, these findings suggest that *FosB*-derived proteins play a crucial role in modulating immediate effector functions in mast cells by limiting mediator storage and reducing sensitivity to antigen-mediated activation and degranulation, without broadly suppressing acute transcriptional responses and independent of FcεRI expression levels. Supporting this, stimulation with 0.1 mM A23187 induced significantly greater and more sustained intracellular Ca²⁺ mobilization in MC^FosB⁻^ BMMCs compared to WT (**Supplementary Figs. 1 C&D**). As A23187 is a calcium ionophore that bypasses FcεRI by directly activating membrane Ca²⁺ channels and releasing Ca²⁺ from endoplasmic reticulum stores^84^, this further supports a role for FOSB/ΔFOSB in regulating intracellular signaling pathways downstream of receptor engagement.

Based on these findings, we next aimed to identify functional targets of FOSB/ΔFOSB—genes that are both differentially bound by FOSB/ΔFOSB in the genome and differentially expressed under baseline conditions or after DNP exposure in IgE-sensitized BMMCs. To achieve this, we integrated the differentially bound peaks identified from FOSB/ΔFOSB CUT&RUN data with differential expression analysis from pre-existing RNA-seq of IgE–Veh-and IgE–DNP-treated BMMCs (**Fig. 3J**).

Differential binding analysis had previously identified over 4,000 genomic regions with significantly altered FOSB/ΔFOSB occupancy after antigen stimulation (**Supplementary Table 3**). Although this analysis is conducted at the level of genomic regions, each peak was subsequently annotated to its nearest gene using HOMER to facilitate biological interpretation. Because multiple peaks can map to the same gene, we collapsed the annotated list to identify 146 unique genes associated with differential FOSB/ΔFOSB binding—hereafter referred to as differentially bound genes (DBGs). Notably, 144 of these 146 DBGs showed increased binding following stimulation, indicating a widespread gain in FOSB/ΔFOSB occupancy. Of these, 36 were also differentially expressed in the RNA-seq dataset, defining a subset of functional FOSB/ΔFOSB target genes with both enhanced DNA binding and transcriptional regulation in response to IgE–antigen stimulation (**Fig. 3K and Supplementary table 6 for full list of genes**). To further validate this dataset and explore potential mechanisms by which FOSB/ΔFOSB modulates mast cell activation, we selected two top-ranked candidate genes for follow-up analysis: the dual-specificity phosphatase Dusp4 and the signaling adaptor Lcp2 (Slp-76). These genes were chosen based on their significant enrichment in both differential binding and expression datasets, as well as their known roles in regulating intracellular signaling pathways in immune cells.

qPCR analysis revealed that DNP stimulation increased *slp-76* expression in both WT and MC^FosB⁻^ BMMCs, although multiple comparisons test revealed that this increase was significant only in WT BMMCs (**Fig. 3L**). In contrast, *dusp4* expression was markedly upregulated only in WT BMMCs (**Fig. 3M**), suggesting that FOSB/ΔFOSB may promote *Dusp4* induction in response to antigenic stimulation. As Dusp4 encodes a nuclear phosphatase that deactivates MAPKs such as ERK1/2, JNK, and p38, thereby attenuating MAPK-dependent signaling cascades^85,86^, this differential expression points to a potential mechanism by which FOSB/ΔFOSB limits mast cell activation—consistent with our finding that IgE-sensitized MC^FosB⁻^ BMMCs exhibit increased ERK phosphorylation.

Next, to determine the physiological relevance of these findings *in vivo*, we next evaluated systemic allergen-like responses in WT and MC^FosB⁻^ mice.

### 3. Mast cell-specific knockout of *FosB* exacerbates histamine release and hypothermic responses in models of systemic anaphylaxis

We first examined whether MC^FosB⁻^ and WT male mice exhibited any basal phenotypic differences. Our analyses revealed no differences in body weight at 6 weeks of age (**Fig. 4A**), locomotor activity, as measured by total distance traveled (**Fig. 4B**) or velocity (**Supplementary Fig. 1E**) in the open field test, or the number of mast cells in mesenteric tissue (**Figs. 4C&D**), indicating that overall physiology remains intact. To confirm that *FosB* deletion was restricted to mast cells, we assessed FOSB and ΔFOSB protein expression in the hippocampus, a brain region where these proteins typically accumulate^51,57,87^. As expected, FOSB and ΔFOSB were detected in WT and MC^FosB⁻^ hippocampal tissue, but they were absent in MC^FosB⁻^ activated BMMCs derived from MC^FosB⁻^ mice (**Fig. 4E**), supporting the cellular specificity of our knockout model.

**Fig. 4.**
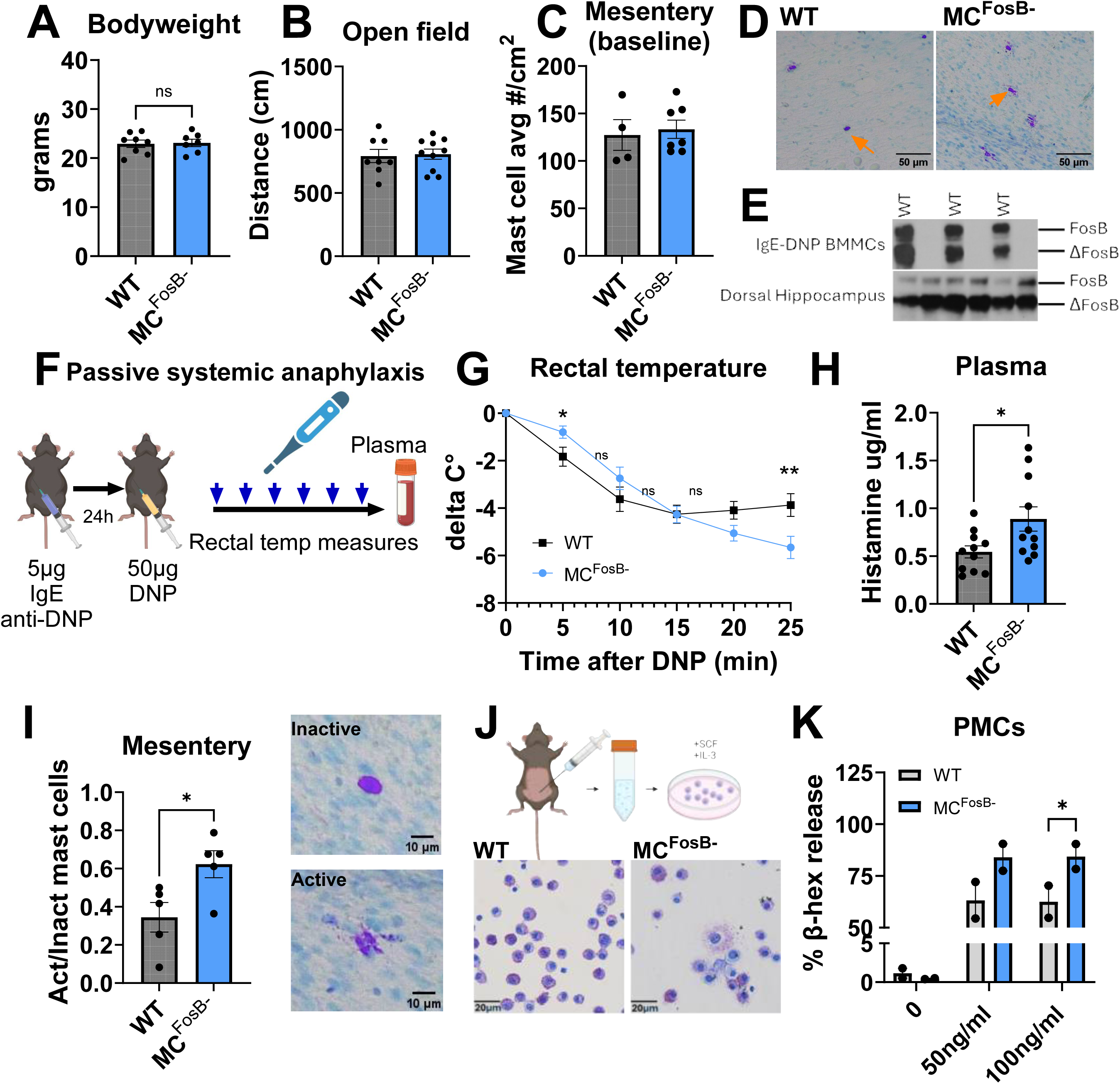
Mast cell-specific knockout of *FosB* exacerbates hypothermic responses, mast cell activation, and histamine release in models of systemic anaphylaxis. A-D. No differences between WT and MC^FosB-^ mice in bodyweight at 6 weeks old (n=8 WT, 7 MC^FosB-^), locomotor activity in the open field test (n=8 WT, 10 MC^FosB-^), or number of mast cells in mesenteric tissues (n=4 WT, 7 MC^FosB-^), **E. Western blot showing FOSB and** ΔFOSB is detected in dorsal hippocampus (bottom) but not BMMCs (top) of MC^FosB-^.F. Timeline showing passive systemic anaphylaxis experiment. **G.** PSA-induced hypothermic responses in WT and MC^FosB-^ mice (n=13 WT, 9 MC^FosB-^, repeated measures two-way ANOVA time x genotype interaction F_(5,100)_ =5.96, p<0.0001; Fisher’s LSD multiple comparisons WT vs MC^FosB-^ at min 5 p=0.04 and min 25 p=0.01). **H.** Plasma histamine 25 minutes after PSA in WT and MC^FosB-^ mice (n=11/group, one-tailed T-test t_(20)_= 2.415, p=0.01). **I.** Active/inactive ratio of mesenteric mast cells in WT and MC^FosB-^ mice (n=5/group, one-tailed T-test t_(8)_= 2.65, p=0.01). **J,K.** Isolation of peritoneal mast cells and IgE-mediated degranulation assessment in WT and MC^FosB-^ mice (n=4/group, two-way ANOVA genotype F_(1,8)_ =11.83, p=0.008; Fisher’s LSD WT vs MC^FosB-^ 50ng/ml DNP p=0.04, 100ng/ml DNP p=0.03).

Next, we used the mast cell-dependent model of IgE-mediated passive systemic anaphylaxis (PSA) to evaluate *in vivo* IgE-FcεRI-dependent mast cell activity. WT and MC^FosB-^ mice were sensitized with an i.p. injection of 5 μg anti-DNP IgE monoclonal antibody and then challenged the following day with 50 μg DNP administered i.p. to induce anaphylaxis, as previously described^82^ (**Fig. 4F**). Rectal temperature was measured every 5 minutes following DNP administration to assess PSA-induced hypothermia, a mast cell-dependent response^88^. 25 minutes after DNP stimulation, mice were euthanized, and blood as well as intestinal mesentery were collected for the assessment of plasma histamine and mast cell histology, respectively. Intestinal mesenteric mast cells are commonly studied in the context of PSA^18^ because they represent a population of tissue-resident mast cells that are highly abundant and strategically located along the gastrointestinal tract, a key site of antigen exposure. Additionally, mesenteric mast cells are well-suited for histological analysis due to their accessibility and high granule density, facilitating the evaluation of degranulation.

Interestingly, we observed a significant interaction between time and genotype in rectal temperature changes following DNP exposure (**Fig. 4G**). At 5 minutes post-DNP exposure, MC^FosB⁻^ mice exhibited a less pronounced drop in rectal temperature compared to WT mice. However, this difference was no longer evident between 10 and 20 minutes, during which both groups displayed similar temperature reductions. By 25 minutes, however, WT mice began to recover from hypothermia (as indicated by an increase in rectal temperature), while MC^FosB⁻^ mice continued to experience a further decline in rectal temperature. Consistent with these results, MC^FosB⁻^ mice showed significantly higher plasma histamine at min 25 (**Fig. 4H**), and an increased ratio of active (degranulating) vs. inactive (non-degranulating) mesentery mast cells (**Fig. 4I).** Because PSA-induced drop in body temperature is dependent on mast cell-derived mediators, including histamine^89^ and chymase^90^, together these results suggest either continued activation of additional mast cells, an overly persistent release of mediators in mast cells from MC^FosB⁻^ mice, or changes to the degradation/elimination of mediators. To test whether tissue-resident mast cells from MC^FosB⁻^ mice show exaggerated mediator release, we cultured mast cells derived from peritoneal cavity **(Fig. 4J)**. Consistent with our *in vitro* and *in vivo* findings, peritoneal MCs from MC^FosB⁻^ mice showed increased degranulation compared to WT **(Fig. 4K)**.

Together, these findings highlight the critical role of FOSB/ΔFOSB in modulating mast cell activity both *in vitro* and *in vivo*, preventing exacerbated inflammatory responses to allergen exposure. However, mast cell functions extend far beyond allergic reactions and include the modulation of immune responses to infection-like stimuli^91–93^. Therefore, we next assessed the effects of MC *FosB* inhibition to LPS-mediated stimulation *in vitro* and *in vivo*.

### 4. Mast cell-specific knockout of *FosB* exacerbates mast cell-dependent hypothermia and increases plasma IL-6 in response to LPS challenge

To test whether *FosB* is involved in infection-like mast cell responses, we stimulated WT BMMCs with 500ng/mL LPS, a component of Gram-negative bacteria^94^ recognized by mast cells through TLR4^95^, and collected the samples 1h later (**Fig. 5A**). Similar to IgE-DNP stimulation, LPS upregulated both *fosB* (**Fig. 5B**) and *ΔfosB* (**Fig. 5C**), suggesting a role for *FosB* in TLR4-mediated mast cell activation. However, despite this transcriptional upregulation, LPS stimulation did not induce detectable FOSB or ΔFOSB protein (**Fig.5D**). This suggests that translation in this context is either diminished or delayed, possibly due to the need for additional signals or a longer course for protein accumulation. This aligns with the fact that TLR-mediated activation does not induce Ca²⁺ mobilization or immediate mast cell degranulation^96,97^. Instead, it primarily triggers the slower synthesis and secretion of cytokines and chemokines, which recruit other immune cells to the site of infection several hours after exposure^95,96,98^.

**Fig. 5.**
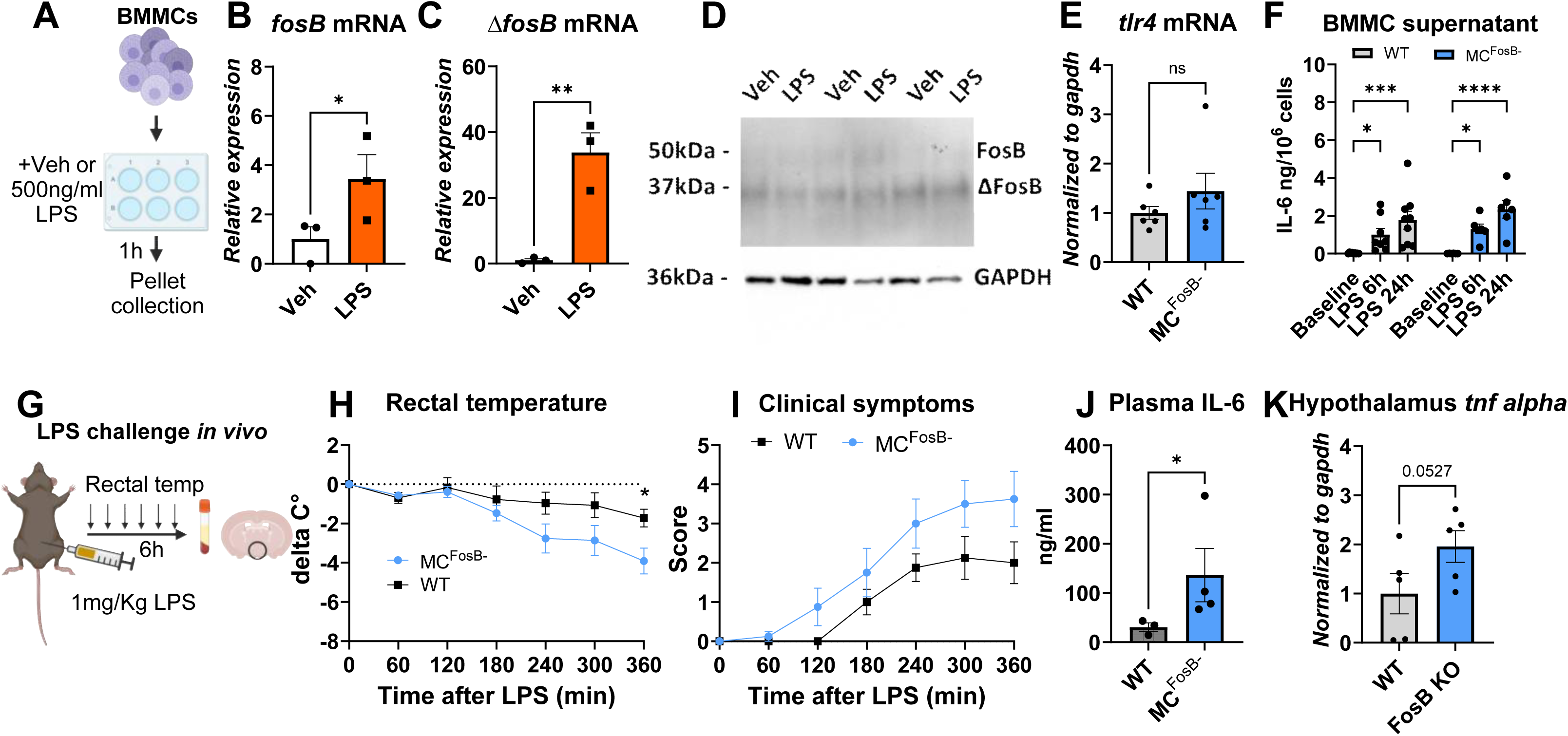
Mast cell-specific knockout of *FosB* exacerbates mast cell-dependent hypothermia and increases plasma IL-6 in response to LPS challenge A. Experimental timeline of BMMC LPS stimulation. **B,C.** Induction of *fosB* (n=3/group one-tailed T-test t_(4)_=2.2, p=0.046, R^2^=0.54) and *ΔfosB* (n=3/group one-tailed t-test t_(4)_=5.47, p=0.002, R^2^=0.88) mRNA in WT BMMCs after LPS stimulation. **D.** Western blot showing no detectable FOSB or **ΔFOSB protein after LPS stimulation. E.** No differences were detected in *tlr4* mRNA expression between WT and MC^FosB-^ BMMCs. **F.** Secretion of IL-6 after LPS stimulation in WT and MC^FosB-^ BMMCs (WT n=9, MC^FosB-^ n=6, two way ANOVA time F_(2,38)_ =19, p<0.0001, Fisher LSD WT baseline vs. 6 and 24h p=0.02 and 0.0002, MC^FosB-^ baseline vs. 6 and 24h p=0.02 <0.0001,respectively). **G**. diagram LPS challenge in vivo. **H.** Rectal temperature measures (n=8/group, repeated measures two-way ANOVA time x genotype interaction F_(6,84)_=3.32, p<0.006, Fisher’s LSD WT vs MC^FosB-^ at hour 6 p=0.02, R^2^=0.35). **I.** Clinical symptoms (n=8/group, repeated measures two-way ANOVA main effect of time F_(2,32)_=30.96, p<0.0001, effect of genotype F_(1,14)_=3.4, p=0.08). **J.** Plasma IL-6 (n=3 WT 4 MC^FosB-^,Mann Whitney test exact p=0.03) in male WT and MC^FosB-^ mice after LPS stimulation *in vivo*. **K**. mRNA expression of hypothalamic *tnf alpha* in male WT and MC^FosB-^ mice after LPS stimulation *in vivo* (n=5/group one-tailed T-test t_(8)_=1.8, p=0.052)

To further assess involvement of *FosB* in this pathway, we measured baseline *tlr4* mRNA levels in WT and MC^FosB⁻^ BMMCs and found no genotype-dependent differences— similar to what we observed for *fcer1* and *fcer2* expression (**Fig.5E**). Next, we measured IL-6 secretion, a key cytokine released in response to TLR4 activation^99^, at 6 h and 24 h post-LPS in both genotypes. Contrary to our hypothesis, IL-6 levels in the supernatant were comparable in WT and MC^FosB⁻^ BMMCs (**Fig. 5F**), suggesting that *FosB* does not significantly regulate IL-6 production downstream of TLR4. However, because mast cell responses to LPS often include the secretion of additional cytokines and chemokines that recruit and activate other immune cells^93,100,101^, we next assessed whether *FosB* inhibition in mast cells could affect physiological responses to LPS *in vivo*.

To investigate this, we stimulated WT and MC^FosB⁻^ mice with 1 mg/kg intraperitoneal LPS and measured rectal temperature every 60 minutes over 6 hours (**Fig. 5G**). This dose of LPS induces mast cell-dependent hypothermia^91^, a response characteristic of severe sepsis^102^. Similar to what we observed in PSA, there was a significant interaction between time and genotype in rectal temperature measures (**Fig. 5H**). By hour 6, MC^FosB⁻^ mice exhibited a more pronounced drop in rectal temperature than did WT mice, along with a trend toward higher clinical scores (**Fig. 5I**), and significantly elevated plasma IL-6 concentrations (**Fig. 5J**). Lastly, given the established role for mast cells in LPS-induced neuroinflammation^92,103^, we measured mRNA levels of the proinflammatory cytokine *Tnfα* in the hypothalamus, and found that MC^FosB⁻^ mice showed a trend toward higher *Tnfα* expression (**Fig. 5K**).

In sum, the observed timeline of body temperature changes and clinical scores, coupled with potentially increased plasma IL-6 levels and hypothalamic *Tnfα*, suggests that mast cells may contribute to exacerbated LPS-induced systemic inflammation in MC^FosB⁻^ mice by enhancing the activation and/or recruitment of other immune cells, thereby amplifying the systemic inflammatory response—possibly independent of mast cell-derived IL-6.

## Discussion

Mast cells play essential roles in physiological homeostasis and immune defense by releasing both pre-stored and newly synthesized mediators in response to diverse stimuli. These effector functions are tightly regulated by changes in gene expression, yet the transcriptional mechanisms that govern mast cell activation remain poorly understood—limiting the development of targeted therapies for mast cell-driven disorders. Although *FosB*, an immediate-early gene involved in stimulus-responsive transcriptional control, is known to modulate functional plasticity in neurons^104^ and microglia^52^, its role in mast cells has remained largely unexplored. Recent reports indicate that *FosB* is widely expressed in human tissue-resident mast cells^1^ and upregulated in murine mast cells in response to glutamate^12^. Building on these observations, we found that both *fosB* and its highly stable splice variant *ΔfosB* are robustly induced in BMMCs following allergy-like IgE–antigen and infection-like LPS stimulations.

Using CUT&RUN profiling in IgE-sensitized BMMCs, we found that FOSB/ΔFOSB bind chromatin under both basal and antigen-stimulated conditions, with antigen exposure leading to increased occupancy and engagement of a partially overlapping but distinct set of genomic regions. This dynamic, stimulus-dependent binding pattern suggests a transcriptional role for FOSB/ΔFOSB in modulating mast cell IgE-antigen mediated activation. To investigate its functional relevance, we generated a mast cell–specific *FosB* knockout mouse (MC^FosB⁻^). BMMCs derived from MC^FosB⁻^ mice displayed increased histamine content and granule abundance at baseline, and upon IgE–antigen stimulation, showed enhanced calcium mobilization, degranulation, and histamine release. *In vivo*, MC^FosB⁻^ mice exhibited exaggerated hypothermia, elevated plasma histamine, and increased activation of tissue-resident mast cells during passive systemic anaphylaxis, indicating a hyperresponsive mast cell phenotype.

To uncover the underlying transcriptional mechanisms, we integrated CUT&RUN and RNA-seq data and identified 36 FOSB/ΔFOSB target genes that were both differentially bound and differentially expressed following IgE-antigen stimulation in BMMCs. Among them, *Dusp4* was selected for follow-up analysis based on known role in negatively regulating MAPK signaling^85,105,106^. qPCR validation revealed that *Dusp4* expression was induced by IgE-antigen stimulation in WT BMMCs but not in MC^FosB⁻^ cells. Given that MAPK pathways contribute to the signaling networks that regulate IgE–antigen–mediated calcium flux and degranulation^107,108^, these results suggest that reduced *Dusp4* expression in MC^FosB⁻^ cells may underlie their heightened reactivity. Together, these findings support a model in which FOSB/ΔFOSB, induced by IgE–antigen stimulation, promote *Dusp4* expression to attenuate MAPK signaling and limit mast cell hyperactivation via a negative feedback loop.

Lastly, building on the observation that FOSB/ΔFOSB limits antigen-driven mast cell activation, we asked whether *FosB* proteins also modulate mast cell involvement in broader immune contexts. Given the role of mast cells in initiating host defense by recruiting and activating other immune cells in response to bacterial pathogens^95,99,109^, we assessed their contribution to inflammation triggered by LPS, a bacterial mimic. While *FosB* inhibition did not alter IL-6 secretion by BMMCs *in vitro*, MC^FosB⁻^ mice exhibited exaggerated mat cell-dependent hypothermia and elevated plasma IL-6 levels following LPS administration. This suggests that FOSB/ΔFOSB may also regulate mast cell contributions to systemic inflammatory responses to infections through indirect or multicellular mechanisms.

### *FosB* proteins are basally expressed in IgE-sensitized BMMCs, where they appear to suppress ERK activation, histamine production, and spontaneous degranulation in the absence of antigen

Our results show that IgE-sensitized BMMCs collected 24 hours after sensitization express both FOSB and ΔFOSB at the protein level, suggesting that these proteins are either constitutively present or persist following IgE engagement. This sustained expression supports a potential regulatory role for *FosB* proteins in modulating mast cell activity under baseline conditions and/or in the period following IgE sensitization.

This is further supported by CUT&RUN data showing that FOSB/ΔFOSB bind to multiple gene regions at baseline in IgE-sensitized BMMCs, and by functional comparisons revealing that unstimulated MC^FosB⁻^ BMMCs exhibit elevated histamine content, increased granule numbers, enhanced ERK phosphorylation, and higher levels of spontaneous degranulation relative to WT BMMCs. GO analysis of FOSB/ΔFOSB-bound regions points to several molecular functions potentially mediating these effects, including terms related to mediator storage, such as amine binding^24,69^, and MAPK signaling^110^, as well as pathways associated with IgE-mediated mast cell activation and proliferation, including MAP kinase activity, interleukin-6 receptor binding, tumor necrosis factor receptor binding, and platelet-derived growth factor receptor binding^7,72–74^. Thus, it is possible that transcriptional regulation of *FosB* proteins in IgE-sensitized mast cells contributes to maintaining a quiescent state and preventing premature activation.

### Under antigen-stimulated conditions, *FosB* proteins appear to further modulate mast cell sensitivity and release of prestored mediators without broadly suppressing acute cytokine synthesis

In addition to baseline differences, MC^FosB^⁻ BMMCs showed increased calcium mobilization and histamine release in response to IgE–antigen stimulation, consistent with an exaggerated effector response. CUT&RUN data integrated with RNA-seq identified several candidate genes potentially underlying these effects, including *Dusp4*, which was validated as a FOSB/ΔFOSB target based on reduced induction in MC^FosB⁻^ compared to WT cells.

*Dusp4* encodes a dual-specificity MAP kinase phosphatase that dephosphorylates ERK, JNK, and p38^111^— key drivers of mast cell degranulation and cytokine production downstream of FcεRI and TLRs^112^. Notably, while MC^FosB⁻^ BMMCs displayed increased MAPK activation and degranulation, they did not differ from WT BMMCs in IL-6 production following IgE–antigen stimulation. This suggests that *FosB* proteins selectively regulate signaling branches controlling degranulation, without broadly altering the transcriptional programs required for cytokine synthesis. One possibility is that the *FosB*–*Dusp4* axis acts with spatial or temporal specificity, attenuating MAPK activity at granule-associated signaling domains rather than at nuclear transcriptional hubs. Alternatively, other MAPK phosphatases may compensate for the loss of *Dusp4* in the regulation of cytokine production, preserving IL-6 output despite heightened ERK signaling.

### Lack of *FosB* in mast cells exacerbates mast cell-dependent physiological responses to allergy-and infection - like stimuli *in vivo*

Using IgE-mediated PSA stimulation *in vivo*, we found that IgE-DNP sensitized MC^Fosb-^ mice, compared to IgE-sensitized WT, showed an exacerbated drop in rectal temperature and release of plasma histamine. PSA-induced hypothermia is a response dependent on mast cells^88^ through two primary mechanisms: first, the release of vasoactive mediators such as histamine, which induce vasodilation and increase vascular permeability^89,113^; and second, the secretion of chymases, a group of proteases that activate TRPV1+ sensory neurons which in turn activate the central thermoregulatory neural circuit leading to a rapid reduction in brown adipose tissue thermogenesis^90^. Although we did not measure chymase release, the fact that chymase is stored together with histamine as a presynthesized mediator in mast cell granules^114^, and given that MC^Fosb-^ mice demonstrate increased degranulation in peritoneal and mesenteric mast cells—which contain high levels of chymase^115^—it is possible that elevated chymases, in addition to elevated levels of histamine, significantly contribute to this exaggerated response in MC^Fosb-^ mice.

We next assessed responses to infection-like stimuli using LPS administration. While LPS stimulation induced expression of both *FosB* and Δ*FosB* in BMMCs, genetic deletion of *FosB* did not alter IL-6 production in vitro, further supporting the hypothesis that *FosB* does not broadly regulate cytokine synthesis in mast cells. However, MC^FosB⁻^ mice, compared to WT, exhibited exaggerated hypothermic responses, more severe clinical symptoms, and trend towards elevated plasma IL-6 secretion following LPS stimulation. Mast cells, which are necessary for LPS-induced hypothermia^91^, also serve as first responders to LPS, recruiting and activating other immune cells such as neutrophils^92,93^. Therefore, these results suggest that *FosB* in mast cells plays a crucial role in limiting the secretion of other mediators beyond IL-6, which could include factors like chemokines^116^ or tryptase^93,117^, to prevent excessive activation of other immune cells involved in LPS-induced systemic inflammation.

### Limitations of this study

This study provides novel insights into the role of FOSB/ΔFOSB in mast cell activation and systemic inflammation. While these findings significantly advance our understanding, several limitations should be noted. First, we were unable to distinguish between the specific contributions of FOSB and ΔFOSB, due to both technical and genetic constraints. Our antibody does not differentiate between the two isoforms—a longstanding limitation in the field—and our mast cell–specific knockout model deletes both isoforms, precluding isoform-specific conclusions.

In addition, although *Dusp4* emerged as a compelling FOSB/ΔFOSB target, other potentially relevant downstream genes remain unexplored. Future work will be needed to map the broader transcriptional network regulated by FOSB/ΔFOSB and determine its relevance in various mast cell contexts. We also did not assess the long-term consequences of FOSB/ΔFOSB induction or its role following repeated stimulation—factors that may be important in chronic or adaptive immune responses. Similarly, *gain-of-function* studies would have provided insight into the sufficiency of FOSB/ΔFOSB in modulating mast cell reactivity, but technical challenges associated with mast cell transfection and transduction currently limit this approach in our lab.

Finally, our experiments were performed exclusively in male mice. Given prior evidence of sex differences in mast cell function^18^, future studies should assess the role of FOSB/ΔFOSB in female mast cells and determine whether sex modulates their impact on inflammatory responses. Overall, this study establishes a foundation for understanding how FOSB/ΔFOSB regulates mast cell activation and provides a strong framework for future investigations into its broader physiological and pathological roles.

## Conclusions

In summary, our research unveils a key role of *FosB* in regulating mast cell responses, highlighting how inhibiting its expression can exaggerate physiological reactions to both allergenic and infection-related stimuli. By exploring the transcriptional landscape modulated by FOSB/ΔFOSB, we have identified potential molecular targets that could refine our understanding and treatment of mast cell disorders. Further, considering recent advances in ΔFOSB research that suggest targeting ΔFOSB in a tissue-or cell-specific manner could be achieved by targeting distinct ΔFOSB isoforms or ΔFOSB-containing multiprotein complexes^104^, this approach emerges as an attractive therapeutic option to explore for disorders associated with exaggerated mast cell activity.

## Acknowledgements

We thank Alicia Withrow and the Center for Advanced Microscopy at Michigan State University for their electron microscopy contributions, and Ken Moon for their excellent technical assistance. This work was supported by a Brain and Behavior Research Foundation 2019 NARSAD Young Investigator Award to NDW; R01 HD072968 to AJM, AJR; R01 AI168014 to AJR, AJM; and R01 DA040621 to AJR.

## Author contributions

NDW, AJM, and AJR conceptualized the study, designed the experiments, and interpreted the data. NDW was primarily responsible for writing the manuscript and performed and analyzed data from most *in vitro* and *in vivo* experiments. BS, MS, and NM assisted with bone marrow cell culture experiments and tissue collection. SY and EN performed and analyzed data from CUT&RUN experiments. DJ performed RNAseq and CUT&RUN data integration analyses. VS and FS assisted with post-mortem tissue analyses. KD assisted with behavioral testing. All authors provided input on the final manuscript.

## Supplemental information

Supplementary methods, Supplementary figure 1 (A-E), Tables S1,S2,S3,S4,S5,S6

## Methods

### Animals

All procedures involving animals were conducted in strict accordance with the ethical guidelines set by the Michigan State University Animal Care Committee. The study utilized in-house bred mice from a cross between MCPT5-^Cre/+^ mice (kindly provided by Dr. Soman Abraham, Duke University, developed by Dr. Axel Roers^80^) and *floxed FosB* mice generated by Eric Nestler’s lab^118^. To generate experimental cohorts, MCPT5-^Cre/+^ homozygous for the *floxed FosB* allele were crossed with MCPT5-^+/+^ mice also homozygous for the *floxed FosB* allele. This breeding strategy yielded approximately 50% of mast cell-specific *FosB*-knockout mice (MCPT5-^Cre/+^; FosB^lox/lox^, referred to as MC^FosB-^), with the remaining 50% serving as littermate controls (MCPT5^+/+^; FosB^lox/lox^, referred to as WT). Genotyping was performed using standard PCR with specific primers: Mcpt5^-Cre^ Forward 5′-ACAGTGGTATTCCCGGGGAGTGT, Mcpt5^-Cre^ Reverse 5′-GTCAGTGCGTTCAAAGGCCA, and FosB loxPu sequence 5′-GCTGAA GGAGATGGGTAA CAG-3′. All animals were housed in same-sex groups of 3-4 per cage, with environmental enrichment including Nestlet and Bed-r’Nest® bedding, maintained at 20–23°C in a controlled 12 h light/dark cycle (lights on 6:00am), and provided *ad libitum* access to food and water. All animals used for experiments in this study were between 6-9 weeks old and only males were used based on previously observed sex differences in mast cell reactivity and mediator release^18^.

### Generation of bone marrow derived mast cells (BMMCs)

Bone marrow progenitor cells were harvested from the femurs of 6-8-week-old male WT and MC^FosB-^ mice. Following established protocols^18,63^, the isolated cells were cultured in a controlled environment at 37°C with 5% CO2. The cultures were maintained in 175 cm² flasks filled with 70 mL of complete medium, which consisted of RPMI 1640 (containing L-glutamine), supplemented with 10% heat-inactivated fetal bovine serum, 1x MEM non-essential amino acids, 10 mmol/L HEPES buffer, 1 mmol/L sodium pyruvate, 100 U/mL penicillin, and 100 µg/mL streptomycin. Additionally, the medium was enriched with recombinant murine interleukin-3 (IL-3; 5 ng/mL) and stem cell factor (SCF; 5 ng/mL) from R&D Systems, Minneapolis, MN. Non-adherent cells were gently transferred to fresh complete medium every 4-5 days. 6 weeks later, purity of mast cell population (>98%) was confirmed using 0.5% toluidine blue staining at pH 0.5 and flow cytometry analysis (BD LSR II) using fluorescently labeled c-kit (BioLegend, San Diego, Calif) and high-affinity IgE receptor (FcεRI) antibodies (eBioscience, San Diego, Calif).

### BMMC stimulations

For IgE-mediated FcεRI crosslinking, BMMCs were sensitized overnight with 0.5 µg/mL mouse monoclonal anti-DNP IgE (SPE-7, Sigma-Aldrich). After sensitization, the media was replaced with fresh complete medium and cells were transferred to a 12-well plate at a density of 2 × 10˄6 cells per well in 1 mL. Following a one-hour acclimation period at 37°C and 5% CO2, either vehicle or 15 ng/mL DNP-HSA (Sigma-Aldrich) was added to each well. Cell pellets and supernatants were collected and immediately flash frozen one hour later for pellet RNA expression measurements and supernatant analyses and after four hours for pellet protein measurements. For LPS stimulation, BMMCs were transferred to a 12-well plate with 2 × 10˄6 cells per well in 1 mL of fresh complete medium. After one hour of incubation at 37°C and 5% CO2, the cells were treated with either vehicle or 500 ng/mL of LPS from Escherichia coli O55:B5 (Sigma-Aldrich). Pellets and supernatants were collected and immediately flash frozen at one hour for pellet RNA expression and supernatant analyses, and at six and twenty-four hours for additional supernatant analyses.

### CUT&RUN

BMMC samples of IgE-sensitized BMMC were incubated with either vehicle or DNP in a 12-well plate for 60 minutes as described above. Following incubation, the samples were centrifuged, and the supernatants were carefully removed. The remaining cell pellets, containing approximately 1×10˄6 cells per sample, were immediately frozen in dry ice. Subsequent procedures including CUT&RUN, library preparation, and sequencing were conducted following previously described methods^66,119^ and are thoroughly described below.

<u>Nuclei preparation:</u> nuclei were extracted from frozen BMMC samples using a Dounce homogenizer to gently break down the tissue in 4 mL of lysis buffer. This buffer was composed of 320 mM sucrose, 5 mM CaCl2, 0.1 mM EDTA, 10 mM Tris-HCl at pH 8.0, 1 mM DTT, 0.1% Triton X-100, 1.5 mM spermidine, and protease inhibitors. The homogenization involved 30 strokes with a loose pestle followed by 30 with a tight pestle. The homogenate was then filtered through a 40 µm strainer. A discontinuous sucrose gradient was prepared by layering 1.8 M sucrose, 10 mM Tris-HCl at pH 8.0, 1 mM DTT, 1.5 mM spermidine, and protease inhibitors under the lysate. Centrifugation was performed at 71,124 RCF for one hour at 4°C, and the resulting nuclear pellet was resuspended in 1 mL of wash buffer containing 20 mM HEPES-NaOH (pH 7.5), 150 mM NaCl, 0.5 mM spermidine, 0.1% Triton X-100, 0.1% Tween-20, 0.1% BSA, and protease inhibitors, all maintained on ice.

<u>CUT&RUN:</u> Nuclei were incubated with BioMag®Plus Concanavalin A (BP531, Bang Laboratories) for 10 min at room temperature. The beads were activated twice with 1.5 ml binding buffer (20 mM HEPES-KOH, pH 7.9, 10 mM KCl, 1 mM CaCl2, and 1 mM MnCl2). Next, the nuclei-bead complex were reclaimed at the magnet and incubated in antibody buffer containing rabbit anti-FOSB (2251, Cell Signaling, 1:50 dilution) antibody at 4°C overnight. The nuclei-beads complexes were then washed with antibody buffer and incubated with antibody buffer containing guinea pig anti-rabbit secondary antibody (ABIN101961, antibodies-online Inc., 1:100 dilution) at 4°C for 1 h. Next, nuclei-bead complexes were washed with antibody buffer and incubated with antibody buffer containing pAGMNase (15-1116, EpiCypher) at 4°C for 1 h. The beads were then washed with wash buffer twice and low-salt buffer (20 mM HEPES-NaOH, pH 7.5, 0.5 mM spermidine, 0.1% Triton X100, 0.1% Tween-20, and protease inhibitor) once before being incubated in calcium incubation buffer (3.5 mM HEPES-NaOH, pH 7.5, 10 mM CaCl2, 0.1% Triton X-100, and 0.1% Tween-20) for 5 min on ice. The beads were reclaimed at the magnet and the reaction was stopped by addition of EGTA-stop buffer (170 mM NaCl, 20 mM EGTA, 0.1% Triton X-100, 0.1% Tween-20, 25 µg/ml RNase, and 20 µg/ml glycogen). DNA fragments were eluted by incubating the samples at 37°C for 30 min with 500 rpm mixing. Next, the samples were centrifuged at 16,000 RCF for 5 min at 4°C and the supernatant was reserved for DNA clean-up using NucleoSpin® Gel and PCR Clean-Up (740609, Takara) based on manufacture instructions.

<u>Library preparation:</u> we used the NEBNext® Ultra™ II DNA Library Prep Kit for Illumina® (E7645, New England BioLabs) with NEBNext® Multiplex Oligos for Illumina® (Dual Index Primers Set 1) (E7600S, New England BioLabs) based on manufacture instructions. Libraries were submitted for sequencing by GENEWIZ on Illumina HiSeq 4000 machine with a 2 × 150-bp paired-end read configuration to a minimum depth of 25 million reads.

<u>CUT&RUN Data Processing and Differential Binding Analysis</u>: Sequencing data were processed using the NGS-Data-Charmer pipeline (GitHub commit date: June 18, 2020; https://github.com/shenlab-sinai/NGS-DataCharmer). Adapter trimming was performed using Trim Galore (v0.6.5). Reads without detectable adapter sequences were secondarily trimmed with Cutadapt (v2.10) to remove 6 base pairs from the 3’ end. Trimmed reads were aligned to the mouse reference genome (mm10) using Bowtie2 (v2.4.1) with the parameters--dovetail--phred33. Duplicate reads were removed using Samtools (v1.10) rmdup.

For visualization, TDF files were generated using IGVtools (v2.5.3), and bigwig files were created using Deeptools (v3.5.0) (bamCoverage--binSize 10--normalizeUsing RPKM). Merged bigwig files were generated from BAM files using Samtools merge and index, followed by normalization with Deeptools (bamCoverage--extendReads--ignoreDuplicates--centerReads--normalizeUsing CPM--binSize 10). Peaks were called for each replicate and its IgG control using MACS2 (v2.2.6)^120^ (macs2 callpeak-f BAMPE--pvalue 0.05--keep-dup) and filtered using IDR (v2.0.4.2)^121^ at a 5% threshold. High-confidence peaks were merged using Bedtools (v2.29.2)^122^. Peak annotation was performed using ChIPseeker (v1.22.1)^123^ in R (v3.6.1). Peak overlap analysis was conducted with HOMER (v4.9) using the mergePeaks module. To compare CUT&RUN binding profiles between IgE–vehicle and IgE–DNP– stimulated BMMCs, differentially bound regions were identified using the DiffBind package (v2.10.0) in R (v3.5.3), applying thresholds of |log₂ fold change| ≥ 1 and FDR < 0.05. Consensus peaksets were derived from reproducible peaks across conditions. Peak regions were annotated to their nearest gene using HOMER^67^ annotatePeaks.pl, and enriched motifs were identified with findMotifGenome.pl using scrambled sequences as background controls.

### Quantitative polymerase chain reaction (qPCR)

BMMC pellets or hypothalamus (collected using a 2 mm diameter punch tool and flash-frozen on dry ice following LPS challenge at bregma ∼0.05–1.05) were homogenized in Trizol (Invitrogen), and total RNA was extracted using the RNeasy Micro Kit (Qiagen #74004). The integrity and concentration of the RNA were assessed using the Nanodrop™ 8000 Spectrophotometer. Subsequently, 200 µg of RNA was converted to cDNA using the Applied Biosystems™ High-Capacity cDNA Reverse Transcription Kit (#4368814). The resulting cDNA was diluted 1:200. qPCR was then employed to analyze changes in mRNA expression (see below for primers used). qPCR reactions were prepared using 10 µL PowerSYBR Green PCR Master Mix (Applied Biosystems #4367659), 2 µL each of forward and reverse primers, 4 µL diluted cDNA, and nuclease-free water to a final volume of 20 µL per well. Thermal cycling was performed with an initial denaturation at 95 °C for 10 minutes, followed by 50 cycles of 95 °C for 15 seconds, 60 °C for 15 seconds, and 72 °C for 15 seconds. Melt curve analysis followed. Each sample was run in duplicate, and relative gene expression was calculated using the ΔΔCt^124^ method, with *Gapdh* as the reference gene.

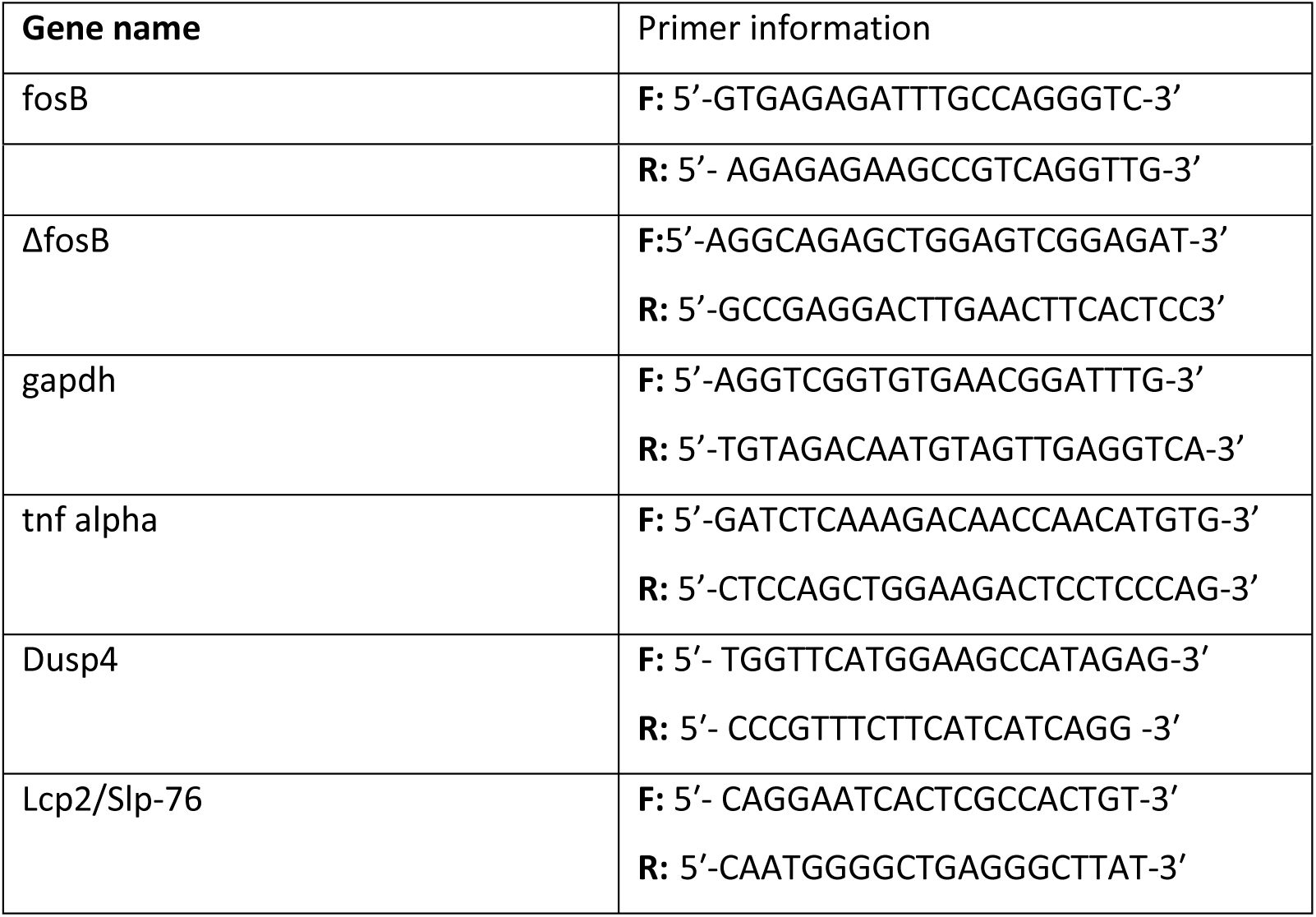

### Western blots in BMMC pellets

Frozen BMMC pellets or dorsal hippocampus sample (collected bilaterally using a 1.5 mm diameter punch tool and flash-frozen on dry ice following LPS challenge at bregma ∼-1.5 to-2.5) were resuspended in 1.5x Laemmli buffer and homogenized through repeated pipetting. 10 µL of each sample was loaded onto a 4-15% SDS-PAGE gel and electrophoresed at approximately 70V for about two hours. Proteins were subsequently transferred to 0.2 µm PVDF Transfer Membranes (Thermo Scientific) at 60V for around 35 minutes. The membranes were incubated overnight with a rabbit anti-FOSB, anti-pERK, or anti-ERK antibodies (#2251, 9101S, and 9102, Cell Signaling Technology) in 5% BSA at 4°C. Following the primary antibody incubation, membranes were washed and incubated with an HRP-conjugated anti-rabbit secondary antibody (PI-1000; Vector Laboratories). Protein signals were detected on film and quantified using ImageJ software. After signal detection, membranes were treated with stripping buffer for 30 minutes and subsequently re-probed with a rabbit anti-GAPDH primary antibody (14C10; Cell Signaling Technology) followed by the same HRP-conjugated secondary antibody. Protein quantification values were normalized to total protein content, as detected by Swift Membrane Stain (G-Biosciences).

### Histamine and IL-6 ELISAS

Histamine levels in BMMC supernatants, BMMC lysates, and plasma were quantified using a competitive histamine ELISA kit (EA31, Oxford Biomedical Research), strictly adhering to the manufacturer’s instructions. IL-6 levels in the supernatant and plasma were determined using a high sensitivity IL-6 ELISA Kit (BMS603HS, Invitrogen) following the manufacturer’s instructions. All assays were conducted in duplicates to maintain consistency and reliability in the results.

### Intracellular calcium mobilization

On the day of the experiment, cells were washed and resuspended in calcium assay buffer. Intracellular calcium changes in response to stimuli were detected using the Fluo 4 NW Calcium Assay Kit (Thermo Fisher Scientific), following the manufacturer’s guidelines. For the measurement of calcium flux, IgE-sensitized cells were stimulated with 0.1 µg/mL DNP/HSA, and unsensitized cells were treated with 0.1 mM A23187. Fluorescence changes were monitored using the FDSS/μCELL kinetic plate reader (Hamamatsu, Japan) at an excitation wavelength of 480 nm and an emission wavelength of 540 nm. This setup provided a dynamic assessment of the cellular response to the added stimuli.

### Electron microscopy

BMMC pellets were fixed with 2.5% glutaraldehyde in phosphate buffer (Electron Microscopy). Samples were post-fixed for 1 h in 1% osmium tetroxide/0.1 M sodium phosphate buffer, dehydrated through a graded series of acetones, and embedded in Spurr epoxy resin. Using a diamond knife, 70-nm sections were cut and mounted on 200-mesh copper grids, then stained with 4% aqueous uranyl acetate for 30 min and Reynolds’ lead citrate for 14min. Samples were observed using a JEOLL 1400 Flash Transmission Electron Microscope operated at 100kV (Japan Electron Optics Laboratory, Japan). The images were taken at magnifications of 500x and 3000x. A blind observer analyzed these images using ImageJ software to quantify whole cell degranulation status and the number of vesicles per cell, respectively.

### RNA-seq Analysis and Differential Gene Expression

Total RNA from IgE–vehicle and IgE–DNP–stimulated BMMCs was sequenced and aligned to the mm10 genome using the same NGS-Data-Charmer pipeline described above. Read counts were imported into DESeq2 (v1.36) in R for differential expression analysis. Genes meeting a significance threshold of |log₂ fold change| ≥ 1.3 and FDR < 0.05 were considered significantly differentially expressed. A more conservative fold-change cutoff was used to limit overestimation of differentially expressed genes in this sensitized cell population.

### Integration of CUT&RUN and RNA-seq

To identify functional targets of FOSB/ΔFOSB, genes with significantly increased binding (from DiffBind) were overlapped with differentially expressed genes from DESeq2 analysis. This allowed identification of genes that were both differentially bound and transcriptionally regulated in response to IgE–antigen stimulation. These overlapping genes were defined as putative FOSB/ΔFOSB target genes.

### Assessment of tissue mast cell numbers and activation status

Small intestinal mesentery windows from WT and MC^Fosb-^ mice at baseline or after passive systemic anaphylaxis were whole mounted on glass slides, fixed with Carnoy’s fixative, and stained with toluidine blue (Sigma-Aldrich, St Louis, Mo) for 30 min, as previously described^63^. Toluidine blue–stained mast cells were counted in 4 nonoverlapping microscopic fields at a magnification of 10x. Each field contained tissue sections that filled the entire high-power field. Mesentery mast cells were counted and scored as inactive (dense staining, non-degranulating) or active (low staining density, granules can be seen outside the cell) by a blind observer using ImageJ in at least 4 mesenteric windows per mouse/treatment. The average number and active/inactive ratio per high-power field was then calculated within each genotype.

### Open field test

Test was performed under red light conditions between 1 and 5 pm (7-11h after lights on) one hour after habituation to the testing room. Mice were placed at the center of a 38 x 38 cm arena and allowed to explore freely for 3 minutes. The velocity and total distance traveled by each mouse was automatically recorded and quantified using Anymaze software (CleverSys).

### *In vivo* passive systemic anaphylaxis

Eight-week-old WT and MC^FosB-^ mice were sensitized with an intraperitoneal (IP) injection of 5 μg of anti-DNP IgE (SPE-7, Sigma-Aldrich) in 100 μL of sterile saline. 24h later, the mice were challenged with an IP injection of 50 μg of DNP-HSA (Sigma-Aldrich) in 100 μL of saline. Rectal temperatures were recorded right before DNP-HAS injection and every 5 minutes for up to 25 minutes following the DNP challenge using a TH-5 Thermalert thermometer (Physitemp, Clifton, NJ), after which animals were euthanized for tissue collection.

### *In vivo* LPS challenge

Eight-week-old WT and MC^FosB-^ mice were challenged with an IP injection of 1mg/Kg of LPS from Escherichia coli O55:B5 (Sigma-Aldrich) in μL of saline. Rectal temperatures were recorded immediately before the LPS injection and subsequently every 60 minutes for up to 360 minutes After the final temperature measurement at 360 minutes, the animals were euthanized for tissue collection. Throughout this period, the mice were closely monitored, and clinical symptoms were assessed and recorded every 60 minutes. The symptoms included reduced activity/alertness, hunched posture, increased respiration rate or labored breathing, ruffled fur, tremors, and inability to move after prodding or to right itself. The overall clinical score for each individual was calculated based on the number of symptoms present, ranging from 0 (no symptoms) to 6 (all symptoms present).

### Statistical analysis and figures

All statistical analyses and data visualizations—except for CUT&RUN and RNA-seq analyses—were performed using GraphPad Prism 9. An alpha level of 0.05 was used to determine statistical significance across all experiments.To compare two conditions (e.g., treatment effects within a genotype or genotype differences under basal conditions), we used one-tailed t-tests or Mann–Whitney tests when data did not meet assumptions of normality. For experiments assessing interactions between genotype and another variable (e.g., treatment), two-way ANOVAs were conducted. In studies involving repeated measures (e.g., calcium imaging, rectal temperature, clinical symptom scoring), we applied repeated-measures ANOVAs with genotype and time as factors.Where significant effects were detected by ANOVA, Fisher’s LSD post hoc test was used for pairwise comparisons. Full statistical results are provided in the corresponding figure legends. In all cases, n represents either an individual mouse or independent cell preparation from a single mouse, as specified in each figure. Data are expressed as mean ± standard error of the mean (SEM). When relevant, effect size correlations (rYl) were calculated using the online effect size calculator https://lbecker.uccs.edu, based on group means and standard deviations. All illustrations and schematics, including the graphical abstract, were created using BioRender™.

## Resource Availability Lead Contact

Further information and requests for resources or reagents should be directed to the lead contact: AJ Robison.

## Materials Availability

This study did not generate any new unique reagents.

## Data and Code Availability

- Raw CUT&RUN sequencing data have been deposited to GEO and are publicly available at: GSE286256.
- Annotated CUT&RUN data tables are included in Supplementary Tables S1–S6.
- This paper does not report original software or code.
- Additional data or materials can be obtained from the lead contact.

**Suuplementary figure 1. A.** Pearson correlation coefficient analysis of FOSB/ΔFOSB coverage among IgE-Veh and IgE-DNP WT and MC^Fosb-^ BMMCs (n=3/group WT, 2/group MCFosb-). **B**. Flow cytometry analysis of BMMCs from WT and MC^Fosb-^ mice. **C,D.** Calcium mobilization in WT and MC^FosB-^ BMMCs in response to A23187 stimulation. Repeated measures two-way ANOVA genotype x time interaction effect F(999, 13986)=6.6 p <0.0001, AUC between seconds 50 and 200 one-tailed T-test t(14)=1.96, p=0.03 R2=0.21) (n=8/group). **E.** WT and MC^Fosb-^ showed no differences in velocity in the open field test.

