## supplementary figure 1 for "FosB/ΔFosB activation in mast cells regulates gene expression to modulate allergic inflammation in male mice"

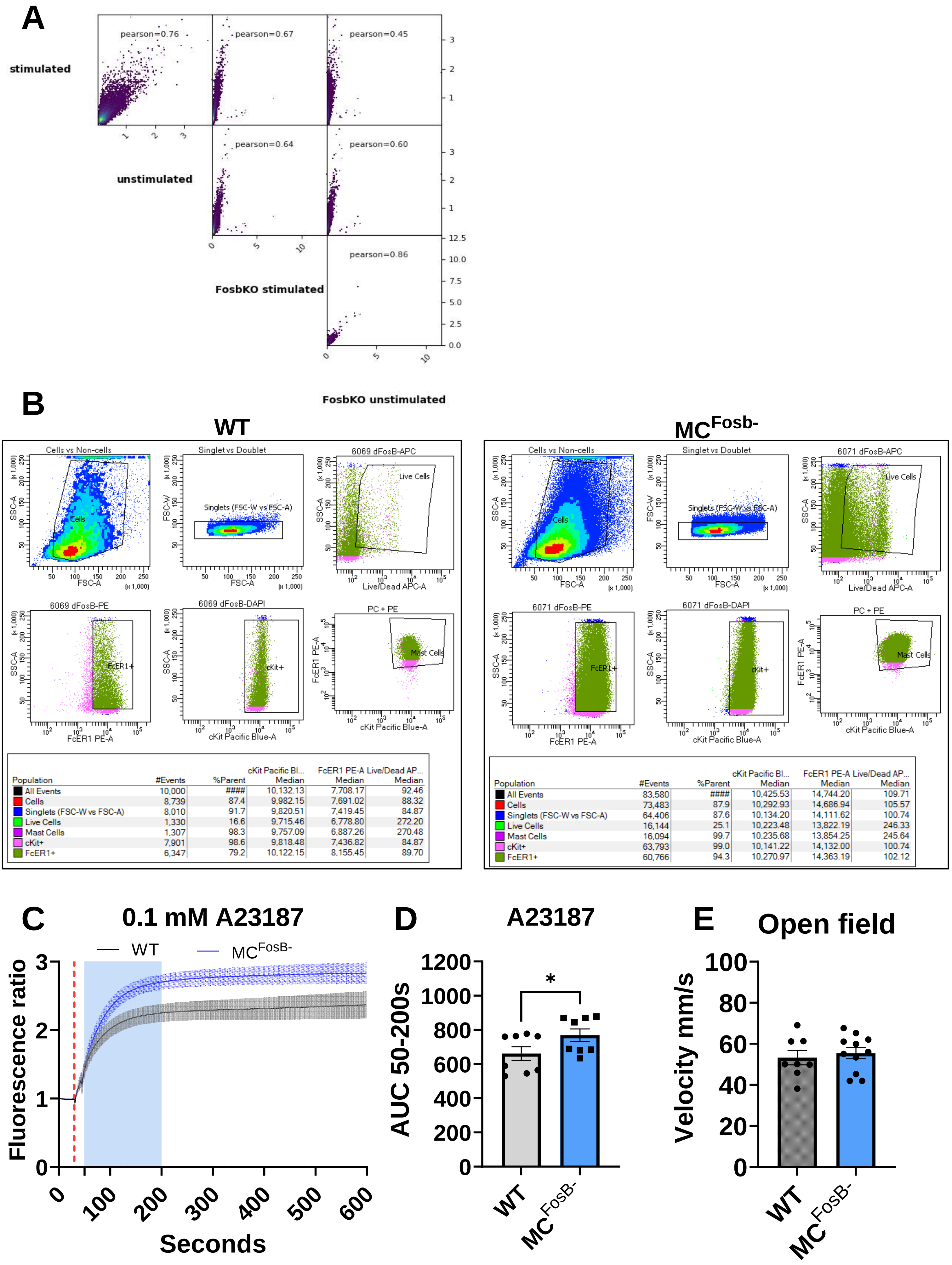


| Population | #Events | %Parent | cKit Pacific Bl... Median | FcER1 PE-A Median | Live/Dead AP... Median |
|---|---|---|---|---|---|
| All Events | 10,000 | #### | 10,132.13 | 7,708.17 | 92.46 |
| Cells | 8,739 | 87.4 | 9,982.15 | 7,691.02 | 88.32 |
| Singlets (FSC-W vs FSC-A) | 8,010 | 91.7 | 9,820.51 | 7,419.45 | 84.87 |
| Live Cells | 1,330 | 16.6 | 9,715.46 | 6,778.80 | 272.20 |
| Mast Cells | 1,307 | 98.3 | 9,757.09 | 6,887.26 | 270.48 |
| cKit+ | 7,901 | 98.6 | 9,818.48 | 7,436.82 | 84.87 |
| FcER1+ | 6,347 | 79.2 | 10,122.15 | 8,155.45 | 89.70 |

| Population | #Events | %Parent | cKit Pacific Bl... Median | FcER1 PE-A Median | Live/Dead AP... Median |
|---|---|---|---|---|---|
| All Events | 83,580 | #### | 10,425.53 | 14,744.20 | 109.71 |
| Cells | 73,483 | 87.9 | 10,292.93 | 14,686.94 | 105.57 |
| Singlets (FSC-W vs FSC-A) | 64,406 | 87.6 | 10,134.20 | 14,111.62 | 100.74 |
| Live Cells | 16,144 | 25.1 | 10,223.48 | 13,822.19 | 246.33 |
| Mast Cells | 16,094 | 99.7 | 10,235.68 | 13,854.25 | 245.64 |
| cKit+ | 63,793 | 99.0 | 10,141.22 | 14,132.00 | 100.74 |
| FcER1+ | 60,766 | 94.3 | 10,270.97 | 14,363.19 | 102.12 |
